# Does red eye fluorescence in marine fish stand out? *In situ* and *in vivo* measurements at two depths

**DOI:** 10.1101/174045

**Authors:** U. K. Harant, F. Wehrberger, T. Griessler, M. G. Meadows, C. M. Champ, N. K. Michiels

**Affiliations:** Department of Animal Evolutionary Ecology, Institution for Evolution and Ecology, Department of Biology, Faculty of Science, University of Tuebingen, Auf der Morgenstelle 28, 72076 Tuebingen, Germany; Saint Francis University, Department of Biology, P.O. Box 600, Loretto, PA, 15940-0600, USA

**Keywords:** *Tripterygion delaisi*, visual contrast, visual communication, coloration

## Abstract

Since the discovery of red fluorescence in fish, much effort has been made to elucidate its potential contribution to vision. However, whatever that function might be, it always implies that the combination of red fluorescence and reflectance of the red iris is sufficient to generate a visual contrast. Here, we present *in vivo* iris radiance measurements of *T. delaisi* under natural light fields at 5 and 20 m depth. We also took substrate radiance measurements of shaded and exposed foraging sites at those depths. To assess the visual contrast that can be generated by the red iris, we then calculated iris brightness in the 600-650 nm “red” waveband relative to substrate radiance. At 20 m depth, *T. delaisi* iris radiance substantially exceeded substrate radiance in the red waveband, regardless of exposure, and despite substrate fluorescence. Given that downwelling light in the 600-650 nm range is negligible at this depth, we can attribute this effect to iris fluorescence. As expected, contrasts were much weaker in 5 m – despite the added contribution of iris reflectance, but we identified specific substrates and conditions under which the pooled radiance caused by red reflectance and fluorescence still exceeded substrate radiance in the same waveband. Due to the negative effect of anesthesia on iris fluorescence these estimates are conservative. We conclude that the requirements to create visual brightness contrasts are fulfilled for a wide range of conditions in the natural environment of *T. delaisi*.

## Introduction

The characteristics of downwelling light changes rapidly with depth in the water column, from directional, bright and spectrally broad near the surface to scattered, dim and spectrally narrow at depth [1-4]. The two main underlying processes are light absorption and scattering [1-4]. Light absorption is particularly strong for longer wavelengths, resulting in a skew towards intermediate, blue-green wavelengths in the visible spectrum. The remaining light is increasingly scattered as it penetrates into the water column resulting in soft, homogeneous lighting that lacks sharp illumination boundaries. These effects have profound consequences for animal coloration as well as visual perception. In shallow water, the ambient spectrum exceeds the visual perception range of fish at both ends of the spectrum. We call this zone the euryspectral zone [5]. With increasing depth, the ambient light quickly narrows down leading into the stenospectral zone, where the spectral range of visual perception can become broader than the available ambient light [5]. Most types of coloration originate from wavelength-specific absorption and reflection by pigments or structural color mechanisms. Possible hues and intensities are therefore strictly limited by their availability in the ambient spectrum. Fluorescent pigments do not have this limitation. They transform absorbed photons of a given wavelength (e.g. in the blue-green range) and re-emit light at longer wavelengths (e.g. yellow or red). Although fluorescent pigments are widespread in benthic marine organisms [6-9], their presence in fish has only recently been confirmed [6, 10-12]. The phylogenetic distribution of red fluorescence in fish correlates with camouflage and sexual signaling [12]. Anthes et al. [12] also found that the presence of conspicuously red fluorescent irides seems to be associated with a micro-predatory lifestyle [5, 13]. Moreover, a recent experimental study indicated that foraging success increases under dim, “fluorescence-friendly” cyan illumination relative to broad spectral illumination at the same brightness in the triplefin *Tripterygion delaisi* [14].

The fluorescence of *T. delaisi* is among the strongest of the fish we have measured thus far [12] and can be perceived by the human eye without the aid of an excitation source or the use of long-pass viewing filters (Figure 1). Yet, it is still weak relative to ambient light. However, visual modelling showed that it is bright enough to generate a brightness contrast between iris radiance and the background radiance that is strong enough to be perceived in conspecifics, at least for non-fluorescent backgrounds [15].

**Figure 1:**
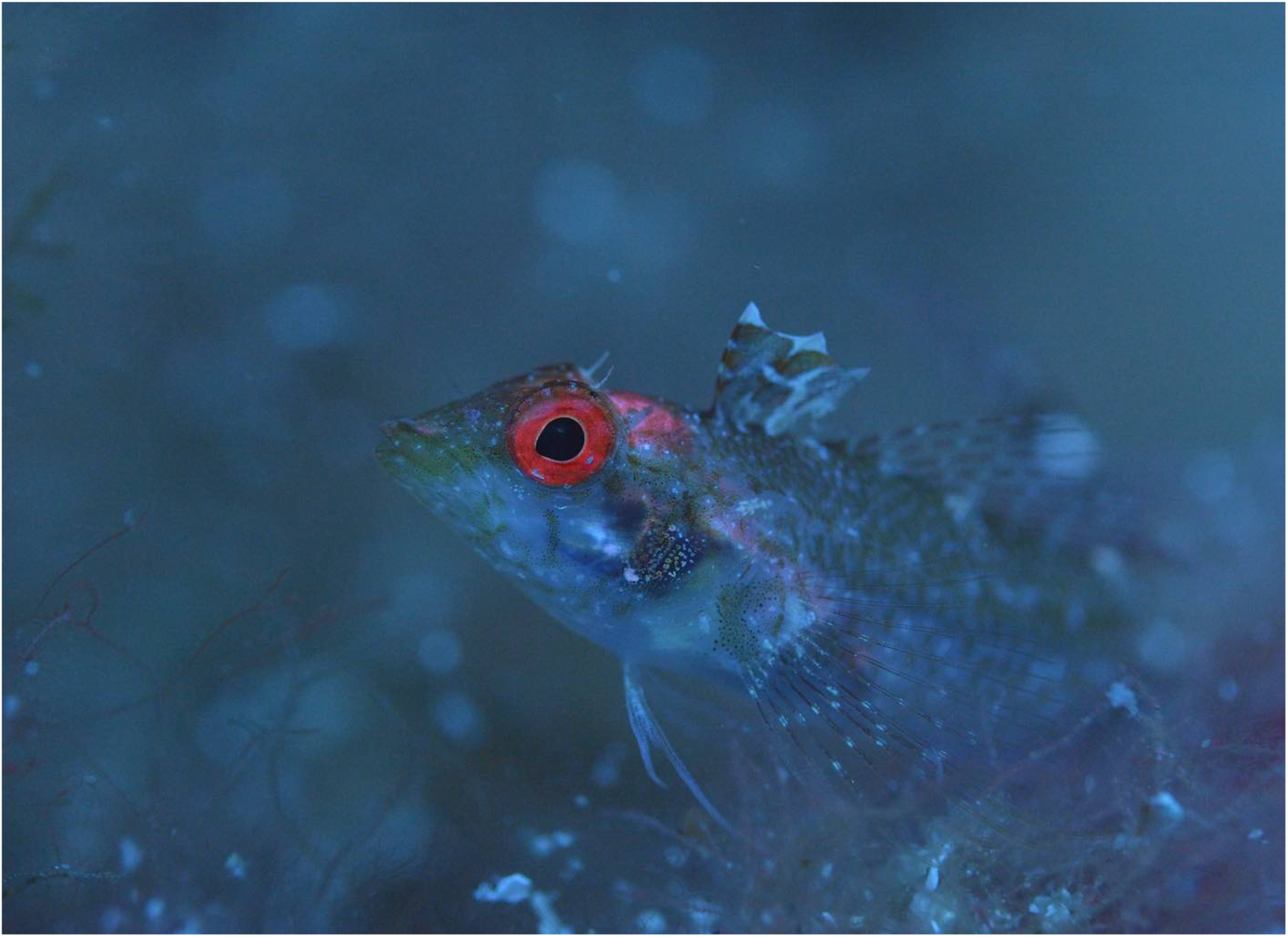
*Tripterygion delaisi* showing conspicuous red iris fluorescence 30 m depth. Picture taken with Nikon D4 + LEE 287 Double C. T. Orange filter and manual white balance, without post-processing (Nico K. Michiels). Note that the LEE filter 287 is not a long pass filter (as is e.g. LEE 105 Orange or LEE 106 Primary Red). It is designed to correct a natural sun lit scene to a warmer spectrum in photography (C. T. = “Correct to Tungsten”). Combined with Manual White Balance, this results in pictures that show colors at depth, including fluorescence, to how they are perceived by a human diver.

Given that natural backgrounds are very diverse, and often fluoresce in the red waveband, we scrutinize the model empirically by directly measuring whether iridal radiance in *T. delaisi* is brighter than the background radiance from the natural substrates on which it lives. To this end, we characterized the natural light environment of *T. delaisi* by measuringthe down- and side-welling light field as well as the radiance of typical substrates under euryspectral (5 m) and stenospectral conditions (20 m). *T. delaisi* uses shaded as well as exposed parts of its home range for foraging, which was also considered in the choice of sites. We also measured iris radiance in anesthetized *T. delaisi in situ* under these conditions. Contrast estimates of substrate and iris radiance allowed us to identify combinations of substrate, depth and exposure under which iris radiance stands out against the background (Figure 1).

## Materials & Methods

The yellow black-faced triplefin *Tripterygion delaisi* is a small, benthic fish from rocky habitats between 5 and 50 m depth along the Mediterranean and eastern Atlantic coasts [16]. It feeds mainly on small, benthic invertebrates [17, 18]. Except for the breeding season, where males develop prominent coloration, individuals are highly cryptic against their natural background, with no obvious sexual differentiation. *T. delaisi* displays highly fluorescent irides with an average peak emission (*λ*_max_) of 609 nm with a full width at half maximum range of 572 nm to 686 nm [15]. Furthermore, it can perceive its own red fluorescence [15, 19], and regulates its fluorescence brightness actively through dispersing and aggregating melanosomes within its melanophores, so that it can switch between near-complete absence of fluorescence to maximum brightness within 10-30 sec [20].

### Field site

Field data were collected at the Station de Recherches Sous-marines et Océanographiques (STARESO) Calvi, Corsica, France in June-July 2014 and 2015. Data were collected while scuba diving at three sites. The shallow site (1) is located just off STARESO and characterized by rocky slopes, steep walls and granite boulders down to 12 m. Exposed hard substrates are covered with a diverse community of green, red and brown algae (Appendix 1). Shaded parts are dominated by coralline red algae and sedentary animals (sponges, cnidarians, bryozoans, ascidians). Flat sandy sediments start at the bottom of the slope and are covered with seagrass (*Posidonia oceanica*), leaving only small patches of rubble and sand. The seagrass meadow slopes gently into deeper water (down to > 30 m). The deep site (2) is located 1 km East of STARESO (“La Bibliothèque”). It features large granite boulders of 1-6 m across from above the surface down to 25 m. A seagrass meadow starts at the bottom of the slope. Areas between the boulders are covered with rubble and sand. The boulders are vegetated mainly by algae including calcareous algae, and some sponges and ascidians, particularly in the permanently shaded parts.

### General spectrometric setup

Radiance measurements were taken with a calibrated PhotoResearch SpectraScan PR-740 radiospectrometer in a custom-made underwater housing (BS Kinetics) with a calibrated MS-75 standard lens. The PR-740 is an all-in-one aim-and-shoot spectrometer with Pritchard optics: It allows to visually focus on a target from a distance with set acceptance angles between 0.1° and 1°. It produces radiance measurements (watts • sr^-1^ • m^-2^ • nm^-1^) in the 380–780 nm range with a 1 nm resolution using a bandwidth of 8 nm. Due to its cooled sensor, this spectrometer captures even very weak signals with little noise at short exposure times. A compass, a level indicator, and an electronic depth gauge were mounted on top of the housing for accurate positioning. During measurements, the dive buddy remained at a safe distance of 5 m in the front of the diver operating the device. Raw data were subsequently corrected for the transmission of the port of the underwater housing and radiance measurements were transformed into photon radiance (photons • s^-1^ • m^-2^ • nm^-1^) by multiplication with wavelength • 5.05 • 10^15^ at each wavelength [21].

### *Radiance of substrates frequented by* T. delaisi

We took spectral measurements throughout the day (07:30 – 18:00) from 29 typical *T. delaisi* sites that were either exposed or shaded at 5 and 20 m depth (Figure 2 A). We defined a substrate to be shaded if it was permanently shaded by e.g. overhanging rocks. Compass direction and surface slope were chosen to cover representative variation. Note that very steep, vertical or overhanging surfaces could not be measured due to handling limitations of the underwater housing, although these areas are also frequented by *T. delaisi*.

**Figure 2:**
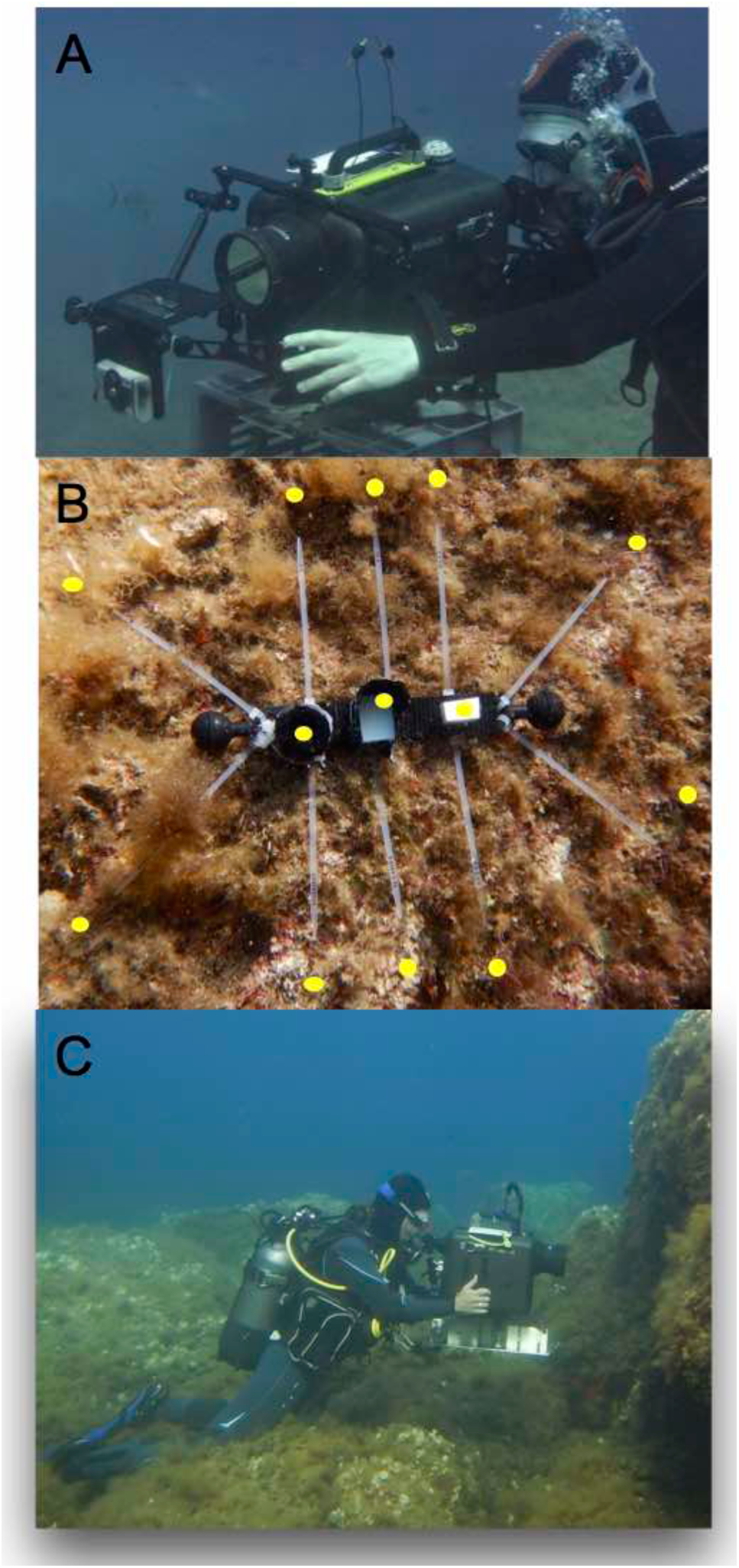
**A:** Substrate radiance measurements were taken at 5 and 20 m depth using a calibrated radiospectrometer (PR740) in a custom made underwater housing (BS Kinetics). **B:** Substrate transect device with reflectance standards in the centre (left to right): black standard, shaded diffuse white standard (PTFE) and non-shaded diffuse white standard (PTFE) (only the last one was used for the calculations presented here). Spectral measurements pointing horizontally onto the substrate were taken approx. 1 cm beyond each of 10 cable binder tips (yellow spot). The length of the central black carrier is 22.5 cm. **C:** Iris radiance measurements taken with a radiospectrometer aiming at a laterally oriented and secured fish at 20 m depth.

To standardize measurements and assess small-scale variation of micro-habitat characteristics, a small transect device was created (Figure 2 B). It defined 10 arbitrary measurement points positioned around three centrally positioned standards: an exposed Polytetrafluorethylen (PTFE) diffuse white reflectance standard (Berghof Fluoroplastic Technology GmbH), (DWS) as a combined measure of downwelling and sidewelling light, a shaded DWS to assess sidewelling light only (not used for any calculations within this study), and a black standard (dark opening of a small vial covered with black cloth inside and outside) as a proxy for the amount of scattered light between spectrometer and substrate. However, the signal of the black standard was mostly too weak to be measured and was therefore not considered for any further calculations. We first measured each standard, then 10 spots on the substrate, each 1 cm above each tip of the 10 measurement pointers (Figure 2 B), followed by a second measurement of each standard. In each transect, all measurements were repeated 3 times, including the standards and the 10 substrate spots. The distance between spectrometer and target was fixed at 60 cm, the minimal focal distance of the spectrometer in the submerged housing. The effect of compass direction was negligible compared to substrate exposure (shaded/exposed) and time of day. We therefore omitted orientation from the results. All raw and derived substrate measurements are provided in Appendix 2 and 3.

To assess whether substrate radiance exceeds the radiance of the DWS as a proxy for the ambient light in the 600–650 nm range, we averaged measurements separately for each specific substrate type within a transect. We then calculated relative radiance as the radiance of that specific substrate type relative to the non-shaded DWS of this transect. Since the non-shaded DWS summarizes the ambient light in a more accurate way (down- plus sidewelling light) compared with the shaded DWS (sidewelling light), we only used the non-shaded DWS for all relative substrate radiance calculations. Values are expected to be smaller than 1, unless substrate fluorescence is strong relative to reflection. Note that we use the term “relative radiance” rather than the more common term “reflectance” because of the combined effects of reflection, transmission (if any) and fluorescence in our radiance measurements.

### Iris measurements of T. delaisi

Iris radiance was measured at 5 m (site 1, *n* = 16 individuals) and 20 m depth (site 2, *n* = 18 individuals) using the same spectrometric setup as described above but with an added SL-0.5x add on macro lens. Additionally, we used a LEE 287 Double C.T. Orange filter, which reduces the abundant blue-green range, allowing longer exposure times to capture better readings in the weak red waveband. We corrected for filter transmission when processing the data (see below). A collection team first caught fish with hand nets at the target depth and brought them to the nearby measurement spot in 50 ml Falcon tubes. The measurement team then anesthetized fish with diluted clove oil and gently placed them in a transparent plastic holder fixed to a small table attached to the front of the spectrometer port (Figure 2 C). The whole head of the fish was fully exposed to the ambient light and the spectrometer. Fish were measured with the side of the eye facing South (sun exposed, more directional light) or North (shaded from direct sunlight, more scattered light). Instead of the Polytetrafluorethylen (PTFE) diffuse white reflectance standard we used waterproof paper (Avery Zweckform) as a diffuse white standard (see Appendix 4 for comparative measurements). The measurement series followed a strict order: First, the white standard was measured, followed by 4 fixed positions within the fluorescent iris (top, right, bottom, left). The measurement angle (shown as a black dot in the viewfinder) was clearly smaller than the width of the iris. Each series ended with an additional measurement of the white standard. Upon completing one eye, the dive buddy turned the fish around for the other eye.

All data were transformed to photon radiance and corrected for reflectance (waterproof paper relative to PFTE, Appendix 2), equipment transmission and the used orange filter as explained above. The measurements taken at the four positions within each eye were averaged per individual. As for the substrate measurements, we express iris radiance as relative iris radiance. All raw and relative radiance measurements are provided in Appendix 5.

### Data analysis

To assess whether iris radiance is stronger than substrate radiance we averaged relative iris radiance as well as relative radiance per substrate type for each condition (2 depths x 2 exposures) for the 600 and 650 nm waveband. We then calculated the Michelson brightness contrast as follows [22]:

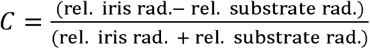

*C* indicates whether iris radiance was stronger (0 < *C* ≤ 1) or weaker (-1 ≤ *C* < 0) than substrate radiance. For graphical representation, we pooled *C* values into 10 categories ranging from < 0 (substrate radiance > iris radiance) to > 0.8. The frequency of cases within each category was then compared between different substrates under the four conditions, and displayed in a mosaic plot. In these plots, each rectangular area is proportional to the abundance of substrate measurements in a particular Michelson contrast category. All Michelson contrasts are provided in Appendix 6.

### Contrast thresholds

Whether a contrast is detectable for fish depends on several factors including the overall brightness in the environment, the size of the stimulus as well as the distance to the stimulus [23]. However, in the euphotic zone, fish with relatively well developed eyes looking at a stimulus roughly matching their size within an ecologically relevant distance have a contrast threshold of 1-2% under bright light conditions [23]. Hence, under optimal daylight conditions, it is assumed that a Michelson contrast between *C =* 0.007–0.05 should be detectable by most fish [24-27].

## Results

### Relative radiance of substrates

At 5 m, relative substrate radiance was largely below one, indicating that fluorescent components in the substrate were too weak to compete with the ambient light (Figure 3). At 20 m, however, relative substrate radiance substantially increased at longer wavelengths, starting at 600 nm and going up to 700 nm (at the borderline of human color vision). At least to some extent, this effect can be attributed to fluorescence from photosynthetic active organisms. Depending on type and exposure, substrate radiation exceeded that of ambient light (indicated by the line at *y* = 1 in Figure 3) by a factor of up to four in the 600–700 nm range.

**Figure 3:**
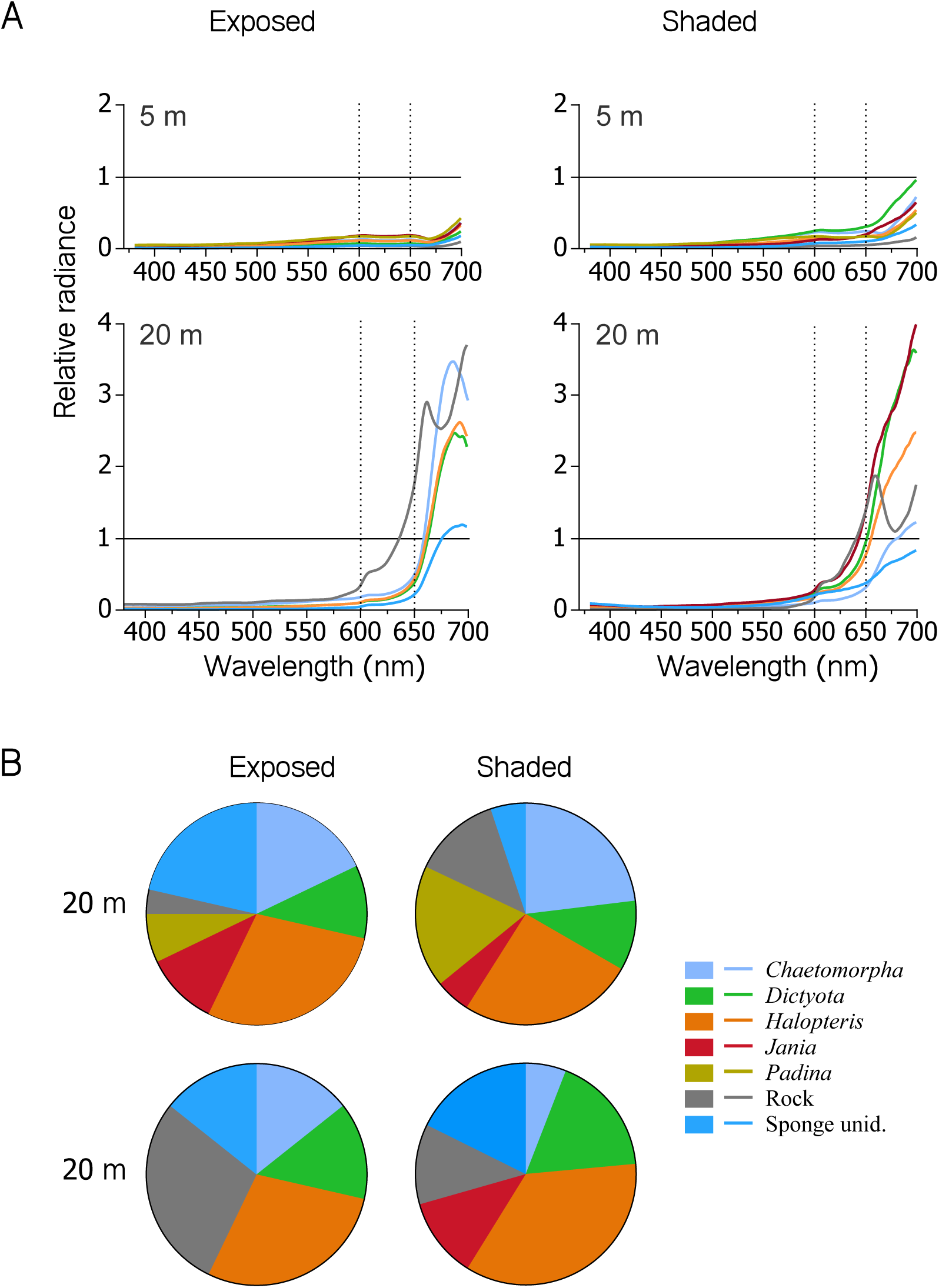
**A**. Line plots showing mean relative radiance of typical *T. delaisi* substrate types as a function of wavelength at 5 and 20 m depth (rows) under sun-exposed and shaded conditions (columns). Values exceeding 1 (black line, referring to diffuse white standard) indicate substrates that emitted more light in that spectral range than was available in the side/downwelling spectrum, a typical signature of strong fluorescence. Dashed lines indicate the waveband of interest (600–650 nm). **B**. Pie charts showing Relative abundance of substrates measured at each combination of depth and exposure. For a detailed species list see Appendix 1.

### Relative radiance of T. delaisi irides

At 5 m, relative radiance of fish irides exceeded 1 in the deep red range (> 680 nm) under shaded conditions (eye facing north) only (Figure 4). This can be explained by the strong red component in the down- and sidewelling light that overrides the fluorescence signal in exposed fish. At 20 m, however, iris radiance exceeded diffuse white standard radiance by up to 9 times (one single measurement), irrespective of exposure – an effect that can only be attributed to iris fluorescence.

**Figure 4:**
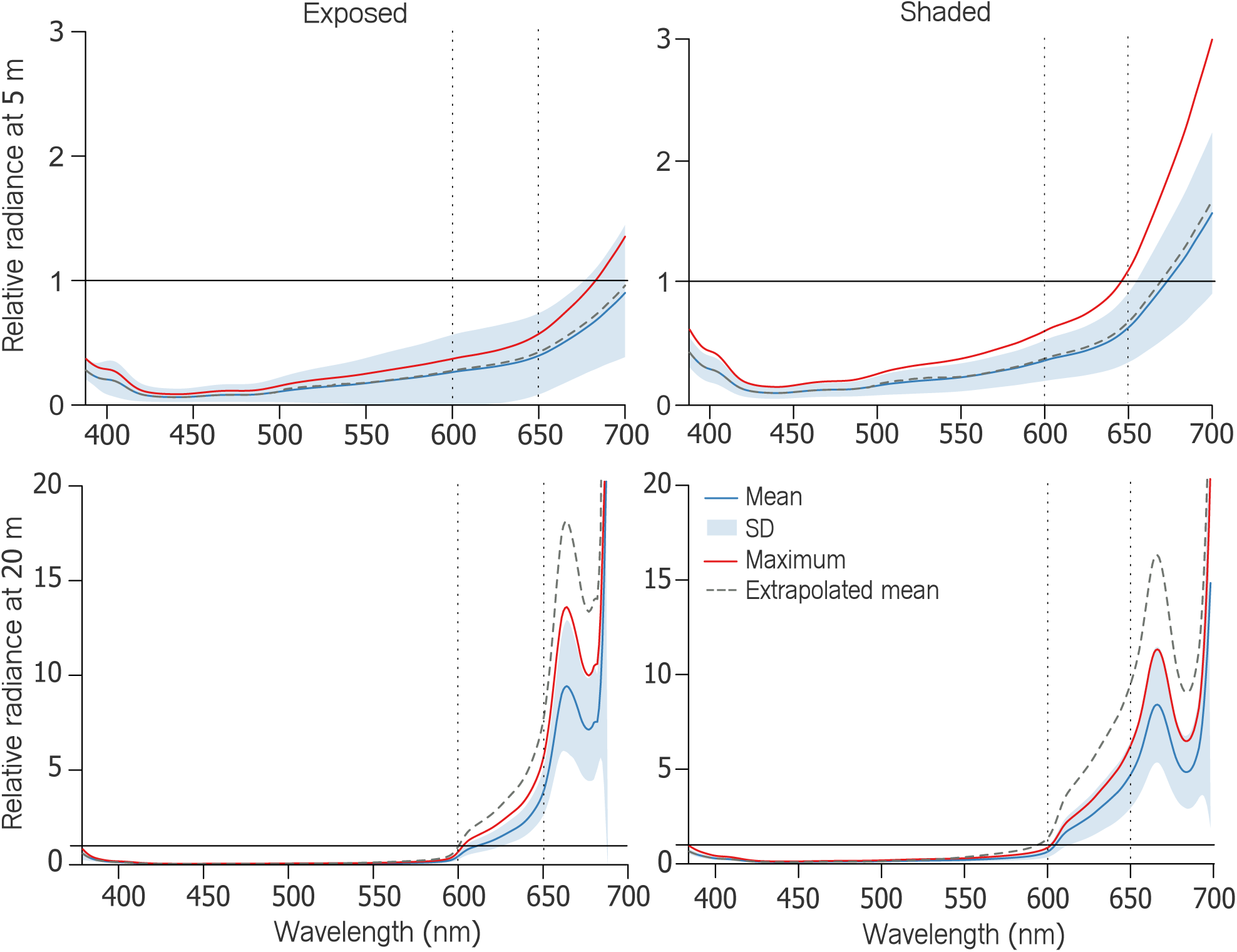
Line plot showing relative iris radiance of *Tripterygion delaisi* as a function of wavelength under exposed (left column) and shaded (right column) conditions at either 5 m (upper row) or 20 m depth (lower row). Blue lines represent means ± SD (shading) of all fish. Red lines indicate the maximum relative radiance averaged across individuals (*n* = 34). Dashed vertical lines indicate the wavelength range of interest (600–650 nm). Values exceeding 1 (horizontal black line) indicate that more photons were emitted by the fish iris at that wavelength than were available in the ambient spectrum, indicative of red fluorescence (assuming absence of specular reflection).

### Comparison between iris and substrate relative radiance

At 5 m, substrate type and exposure determined whether iris radiance exceeded substrate radiance (Figure 5): More contrast prevailed under shaded conditions. Under exposed conditions, iris radiances exceeding substrate radiance were limited to bare rock and sponge substrates, as these two exhibit distinct fluorescence compared to others. At 20 m, however, iris radiance was always stronger in the target wavelength range regardless of substrate type and exposure (Figure 5). The time of the day affected iris contrast only at 5 m depth. Under exposed conditions, iris radiance is more likely to exceed substrate radiance in the morning than in the afternoon (Figure 6). Conversely, under shaded conditions, iris radiance always exceeded substrate radiance in the afternoon, but less so in the morning. An effect of the time of the day was absent at 20 m (data not shown).

**Figure 5:**
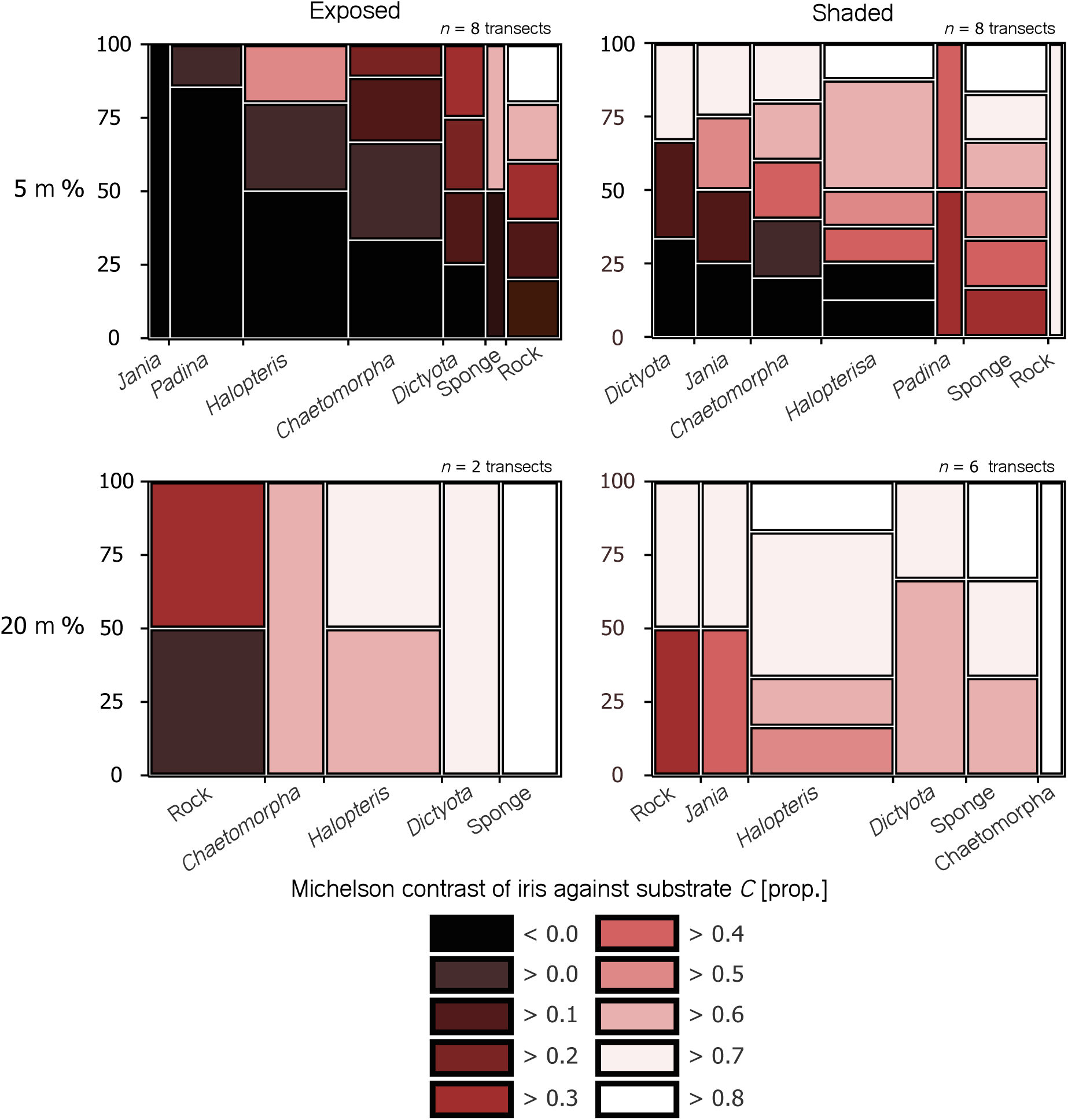
Mosaic plot showing the relative distribution of Michelson contrasts in the target waveband (600–650 nm) (Y-axis) within the 8 commonest substrates (X-axis) at 5 and 20 m depth under exposed or shaded conditions. We defined 10 Michelson contrast categories, where all except the darkest (black) shading indicate iris radiances exceeding substrate radiance. Substrates were ranked from the lowest to the highest brightness contrast.

**Figure 6:**
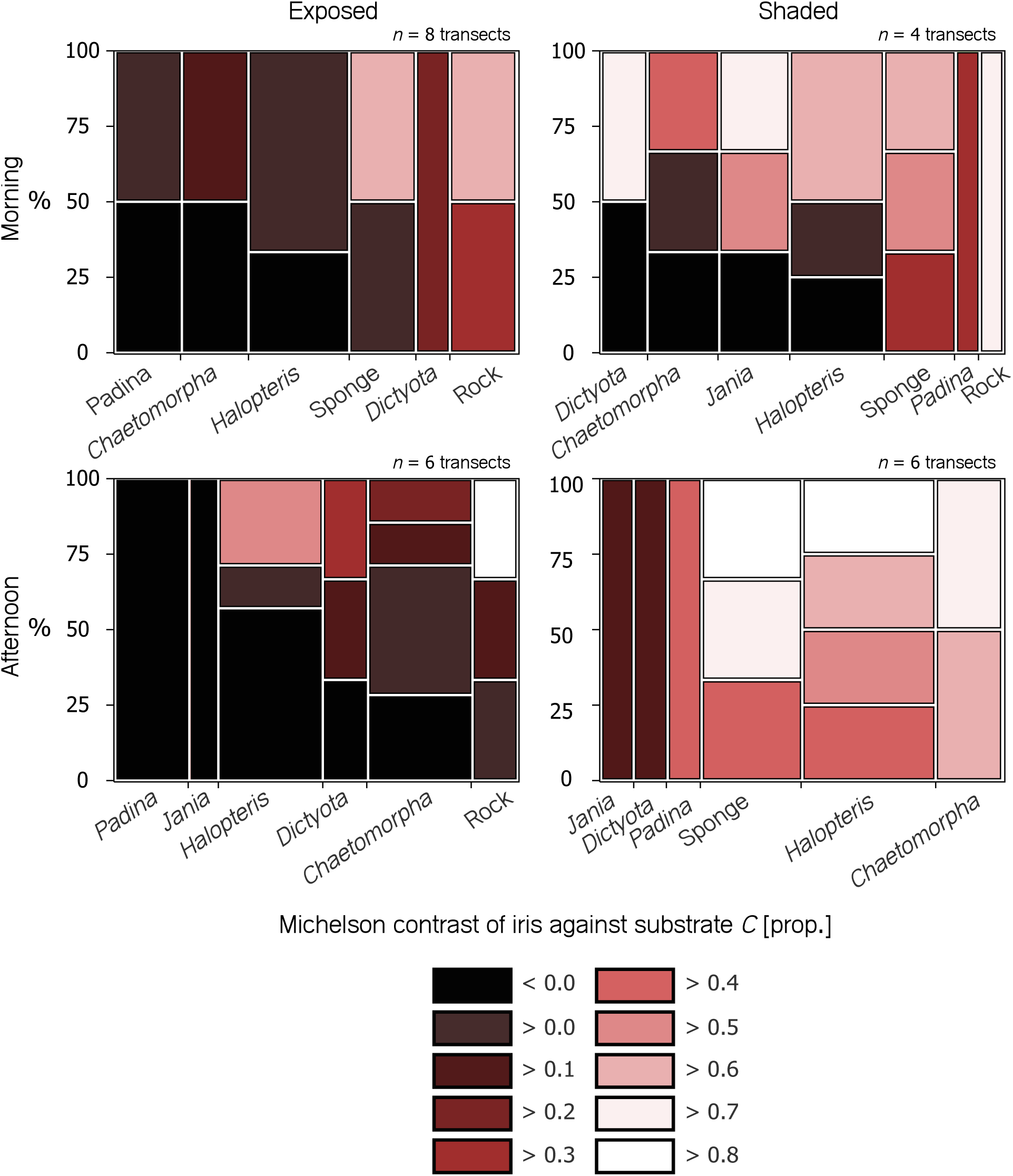
Mosaic plot showing the relative distribution of Michelson contrasts in the target waveband (600–650 nm) (Y-axis) within the 8 commonest substrates (X-axis) at 5 and 20 m depth in the morning (06:00 – 11:30, top) or afternoon (12:00 – 18:00, bottom) under exposed (left) and shaded (right) conditions. Values > 0 (dark red to white) are cases where iris radiance exceeds substrate radiance in the relevant wavelength range. Substrates were ranked from the lowest to the highest brightness contrast.

### Anesthesia effect

Using clove oil for anesthesia leads to a noticeable reduction in iris radiance due to expanding iridal melanophores [20]. This is especially true for fish from 20 m depth, where anesthesia decreases iris radiance by 46 % on average compared with non-anesthetized fish. Fish caught at 5 m depth reduced their iris radiance by only 14 % on average after being anesthetized. The depth-dependency can be explained by reduced iridal melanophore densities in individuals at depth [20, 24]. Therefore, and conservative regarding our research hypothesis, all measurements presented here underestimate natural iris radiance, particularly in individuals from deeper water (see estimated mean relative iris radiance in Figure 4).

## Discussion

Iris radiance of *Tripterygion delaisi* in the 600–650 nm wavelength range exceeded that of the available substrates under stenospectral conditions at 20 m, irrespective of substrate type, exposure and time of day. Under euryspectral conditions at 5 m, however, iris radiance was often less bright compared with the reflection of the stronger red component in the ambient light. Yet, even at this depth, iris radiance exceeded substrate radiance in shaded sites dominated by side-welling blue-green scatter. Due to the effect of anesthesia on iris fluorescence, these estimates are conservative. Consequently, our work confirms empirically that iris radiance (reflectance + fluorescence) in *T. delaisi* is strong enough to generate visual brightness contrasts in a large part of its natural environment, particularly at deeper sites [5, 24]. Bitton et al. [15] produced similar results through modelling, but assuming an achromatic, non-fluorescent background. Our results now confirm that those results may hold against complex, partly fluorescent backgrounds as well. The lack of longer wavelengths along with the reduced overall brightness make stenospectral habitats particularly suitable for the use of fluorescence to generate contrast [5, 24]. This might explain why some particularly strongly fluorescing species are restricted to deeper water such as several species of *Bryaninops*, *Ctenogobiops*, or *Crenilabrius* [12]. Although Anthes et al [12] did not find a correlation between increasing depth and red fluorescence across species, it is safe to assume that red fluorescence is more likely to contribute to vision in deeper water rather than in shallow water. In fact, when analyzing individuals collected at 5 and 20 m within single species (including *T. delaisi.*), Meadows et al [5] found that fluorescence brightness increased with depth when measured under identical laboratory conditions. Although we did not investigate the functionality of red fluorescence, our results are nevertheless in line with previous suggestions that intraspecific communication [15] or even prey detection using active photolocation might be facilitated through red fluorescence [12, 14].

### Limitations of measuring different T. delaisi habitat types

Although we identified several substrate types on which red fluorescence is particularly likely to generate perceptible brightness contrasts, we need to emphasize that certain typical microhabitats could not be measured. Due to handling limitations of the underwater housing, and the need for upward facing substrates to place the transect device (Figure 2 B), we could not take measurements from underneath overhangs or in crevices, which are also important for triplefins. However, given that these shaded sites are exclusively illuminated by blue-green, side-welling light, relative iris radiance in the long-wavelength range should be high, except where encrusting red calcareous algae are common. The latter often cover large areas inside crevices and exhibit very strong red fluorescence.

### Conclusions

We found that in *T. delaisi*, iris radiance in the 600-650 nm bandwidth exceeds the radiance of all measured natural backgrounds in deeper water. This effect can largely be attributed to red fluorescence, which strongly exceeds reflection at depth. But even in shallow water, where red reflectance is considerable [15], iris radiance exceeded that of the background for several substrate types, particularly when shaded. Our findings show that iris radiance can generate relevant visual brightness contrasts against its natural background and might therefore also be relevant in terms of prey detection or intra-specific communication.

## Acknowledgement

We would like to thank Andreas ‘Oeli’ Oelkrug, Christopher Rader, and Gregor Schulte for technical assistance. Additionally, we want to thank our hosts in STARESO, Corsica, for providing excellent working conditions both below and above the surface.

## Authors’ contributions

UKH and NKM designed the experiments and optimized the methodology. Data collection: UKH, NKM, MGM, CMC, FW, TG. Data analyses and drafting of the manuscript: UKH. Editing of the manuscript: UKH, NKM, MGM, FW, TG, CMC. All authors read and approved the final manuscript.

## Author details

Animal Evolutionary Ecology, Institution for Evolution and Ecology, Department of Biology, Faculty of Science, University of Tuebingen, Auf der Morgenstelle 28, 72076 Tuebingen, Germany. This work was funded by a Reinhart Koselleck Project Grant Mi482/13-1 from the Deutsche Forschungsgemeinschaft to N.K.M.

---

**Appendix 1:** Species list and photographical documentation of measured substrates.

**Appendix 2:** Raw data of all substrate measurements taken.

**Appendix 3:** Relative radiance data of substrate measurements.

**Appendix 4:** Comparison between diffuse white standards (PTFE vs. underwater proof paper)

**Appendix 5:** Raw and relative radiance data of *in situ* iris measurements taken in *T. delaisi.*

**Appendix 6:** Michelson contrast calculations of iris against substrate.

